# Multi-species cryoEM calibration and workflow verification standard

**DOI:** 10.1101/2024.08.05.606612

**Authors:** Daija Bobe, Mykhailo Kopylov, Jessalyn Miller, Aaron P. Owji, Edward T. Eng

## Abstract

Cryogenic electron microscopy (cryoEM) is a rapidly growing structural biology modality that has been successful in revealing molecular details of biological systems. However, unlike established biophysical and analytical techniques with calibration standards, cryoEM has lacked comprehensive biological test samples. We introduce a cryoEM calibration sample that is a mixture of compatible macromolecules that can be used not only for resolution optimization but also provides multiple reference points for evaluating instrument performance, data quality, and image processing workflows in a single experiment. This combined test specimen provides researchers a reference point for validating their cryoEM pipeline, benchmarking their methodologies, and testing new algorithms.

## Introduction

Cryogenic electron microscopy (cryoEM) has grown increasingly popular in structural biology as the quality and reliability of cryo-transmission electron microscopes have improved. However, due to the complex nature of the instrumentation, each microscope in operation is unique. Microscope builds can feature different gun sources, accelerating voltages, condenser systems, aberration correctors and filters, and camera types. Each microscope is affected by its local environment such as temperature, humidity, vibrations and electromagnetic fields, all of which influence data quality ^1,2^. To ensure optimal performance, cryoEM practitioners rely on workflow validation tests using an analytical benchmark standard ^3–5^.

Current requirements for cryoEM benchmarking standard are:

- **Accessible**: can either be commercially purchased or has straightforward sample preparation with low maintenance requirements.
- **Stable**: can be stored for prolonged periods of time without compromising structural integrity.
- **Homogeneous**: minimal conformational and compositional heterogeneity.
- **Reproducible**: grid preparation can be standardized reducing grid-to-grid variability.

Common benchmarking samples, including ribosomes, tobacco mosaic virus, beta-galactosidase, aldolase and apoferritin, meet the above criteria and have proven useful for resolution optimization ^3,6–8^. Despite its importance, maximizing resolution is not the only goal of the cryoEM workflow. The datasets acquired for benchmarking resolution can also be used to support other aspects of the cryoEM workflow such as pixel size calibration, neural network training, helical and single-particle processing, etc.

Here, we introduce a dedicated cryoEM calibration standard called the “EM ladder”, which extends beyond resolution benchmarking. A key aspect of the EM ladder is its ability to be used reproducibly under a wide variety of experimental conditions and pixel sizes for calibration, commissioning, and certification of standard operating procedures. Similar to calibration standards for SDS-PAGE, size exclusion chromatography, and mass photometry, there is not a single calibration reagent, but a mixture of components to be compatible with the full experimental workflow. We have combined a mixture of four samples: apoferritin (ApoF - O symmetry, ∼486 kDa, 20 kDa monomer), beta-galactosidase (β-gal - D2 symmetry, ∼465 kDa, 116 kDa monomer), a virus-like particle (PP7 - I symmetry, ∼3.4 MDa, 28 kDa monomer), and tobacco mosaic virus (TMV - H symmetry ∼40 MDa, 18 kDa monomer).

The main focus of this calibration standard is to provide a stable and well-characterized reference point that ensures that the full workflow of grid preparation, data collection, processing, and analysis is reliable. As cryoEM continues to evolve and be applied to new fields, this calibration standard broadens the scope where structural insights are not just high in resolution but also high in accuracy, enabling the full potential of structural biological research.

## Materials and Methods

### Samples

The stock apoferritin (ferritin human H chain, ApoF) was purified in 50 mM Tris pH 7.5, 100 mM NaCl, and 0.5 mM TCEP by the Center on Membrane Protein Production and Analysis (COMPPÅ) at the New York Structural Biology Center. The original plasmid LF2422 contains ferritin human H chain cloned into pGEX2T with a TEV site instead of thrombin from the Protex facility at the University of Leicester, and gifted to NYSBC by Louise Fairall and Christos Savva.

Thyroglobulin (ThG) was purchased from Sigma (Product Number T1001). The lyophilized powder (40 mg) was reconstituted in a storage buffer consisting of 20mM HEPES pH 7.4, 150 mM NaCl buffer.

Beta-galactosidase (β-gal) was purchased from Sigma (Product Number G5635). The lyophilized β-gal (50 mg) was reconstituted in a storage buffer consisting of 50 mM Tris HCl, 10 mM MgCl2, 10 mM beta-mercaptoethanol at pH 7.3. PP7 VLPs were a gift from M.G. Finn’s group. PP7 WT sample was provided as a 1 mg/mL solution in 100 mM PBS pH 7.0 as previously described ^9^.

Tobacco Mosaic Virus (TMV) at a stock concentration of 34.85 mg/mL in TBS was a gift from Ruben Diaz-Avalos.

For the final mix that was used for cryoEM imaging, each protein was diluted with 50 mM HEPES pH 7.5, 100 mM NaCl. This buffer was chosen for its compatibility with each protein. The final concentration for each protein prior to mixing was as follows: 0.16 mg/mL apoferritin, 0.10 mg/mL PP7, 1 mg/mL β-gal, 0.17 mg/mL TMV. To create the final mix, 2 μL of each protein was added together. The concentrations of each protein in the final mix were as follows: 0.04 mg/mL ApoF, 0.025 mg/mL PP7, 0.25 mg/mL β-gal, 0.0425 mg/mL TMV. The mix was aliquoted, snap frozen, and stored at –80C for later use.

### Negative stain grid preparation

Continuous carbon grids made in-house were plasma cleaned with a hydrogen/oxygen mix for 30 s on a Gatan Solarus. Two 20 μL droplets of distilled water were added to parafilm followed by three 20 μL droplets of 2% uranyl formate. 3 μL of sample was applied to the continuous carbon grid for 45 s to 1 min. The grid was side blotted, then followed the sequence of dipping into a droplet carbon side down and side blotting for both water droplets and two uranyl formate droplets. The grid was held in the last uranyl formate droplet for 1 min before side blotting and back blotting to remove excess stain.

### CryoEM grid preparation

3 μL of freshly thawed protein from the final mix was applied to plasma-cleaned UltrAuFoil R1.2/1.3 300 mesh holey gold grids (Quantifoil® Micro Tools, Großlöbichau, Germany), blotted for 2.5 s after a 30 s wait time, and then plunge frozen in liquid ethane, cooled by liquid nitrogen, using the Vitrobot Mark III (FEI, Hillsboro, OR) at 75% relative humidity.

### Screening

For screening a Thermo Fisher Scientific Tecnai 12 with a Gatan TVIPS F416 CMOS camera was operated at 120 kV with a 100 μm objective aperture. Images were collected at a pixel size of 2.46 Å using Leginon ^**10**^ at 800 ms per exposure with a dose of ∼50 e-/Å2 and at a nominal defocus range of 2–4 μm.

### Data acquisition

For final data acquisition a Thermo Fisher Scientific Titan Krios G2 with a spherical aberration corrector and a post-column Gatan Image Filter (GIF) and Gatan K2 Summit was operated at 300 kV with a 70 μm C2 aperture and 100 μm objective aperture. Images were collected in counting mode with a 30 eV slit width and calibrated pixel size of 1.096 Å using Leginon ^**10**^ at a dose rate of 6.95 e−/Å^2^/s with a total exposure of 10 s, for an accumulated dose of 69.46 e−/Å^2^. A total of 3,996 images were collected at a nominal defocus range of 1.5–2.5 μm. Given data retention policies the original movies were not archived and the MotionCor2 with dose weighting ^**11**^ aligned summed images by the Appion ^**12**^ pre-processing pipeline were stored as JPEG for Appion image viewer functionality ^**13**^. For this study, those images were converted back to 32-bit MRC files using EMAN2 ^**14**^ for further processing in cryoSPARC ^**15**^.

### Image Processing

The 19jan04d dataset was manually curated and 1,862 out of the 3,996 images had three or more protein types. After conversion to 32-bit MRC, selected images were imported into cryoSPARC as micrographs for CTF estimation, particle picking and extraction, and subjected to 2D and 3D classification, initial model generation and refinement. Processing settings for the one-shot processing are reported in Table 1 with all non-default settings highlighted. Additional details and intermediate results of the processing of individual components are provided in supplemental information.

**Table 1.** CryoSPARC processing parameters for one-shot parallel processing. Gray rows - processing common to all four species; blue rows - ApoF specific steps; red rows - β-gal specific steps; yellow rows - PP7 specific steps; green rows - TMV specific steps.

| # | Job type | Input | Non-default settings | Outcome |
| --- | --- | --- | --- | --- |
| 1 | Import micrographs | Dose-weighted micrographs | Pixel size: 1.096<br>Accelerating Voltage: 300<br>Spherical aberration: 0.01<br>Total exposure dose: 70 | 1,862 micrographs imported |
| 2 | Patch CTF | Imported micrographs | - | 1,862 micrographs CTF-estimated |
| 3 | Blob picker | CTF-estimated micrographs | Min. particle diameter: 110<br>Max. particle diameter: 140<br>Use ring blob: true<br>Mics to process: 100<br>Maximum number of local maxima to consider: 100 | 43,712 picks |
| 4 | Extract micrographs | 43,712 blob picks | Extraction box size: 512<br>Fourier crop to box size: 256 | 31,835 particles |
| 5 | 2D classification | 31,835 particles | - | 50 particle classes and 50 templates |
|  | Select 2D | 50 templates | - | 4 templates selected (Figure 2A) |
| 6 | Template picking | CTF-estimated micrographs, four selected templates | Particle diameter: 420<br>Min. separation dist: 0.4 | 343,918 picks |
|  |  |  | Maximum number of local maxima to consider: 1000 |  |
| 7 | Extract micrographs | 343,918 blob picks | Extraction box size: 480 | 262,074 particles |
| 8 | 2D classification | 262,074 particles | - | 50 particle classes and 50 templates |
| 9 | Select 2D | Output of step 8 | - | ApoF: 1 class - 101,354 particles |
| 10 | ApoF *ab-initio* reconstruction | Output of #9 | - | Initial model |
| 11 | Homogeneous refinement: ApoF | Particles and initial model from #10 | symmetry "O" | Final map |
| 12 | Select 2D | Output of step 8 | - | β-gal: 3 classes - 7,606 particles |
| 13 | β-gal *ab-initio* reconstruction | Output of #12 | - | Initial model |
| 14 | Homogeneous refinement: β-gal | Particles and initial model from #13 | symmetry "D2" | Final map |
| 15 | Select 2D | Output of step 8 | - | VLP: 1 class - 2,351 particles |
| 16 | PP7 *ab-initio* reconstruction | Output of #15 | - | Initial model |
| 17 | Homogeneous refinement: PP7 | Particles and initial model from #16 | symmetry "I" | Final map |
| 18 | Select 2D | Output of step 8 | - | TMV: 1 class - 4,977 segments |
| 19 | 2D classification (straightening TMV 2D classes) | 4,977 segments from TMV selected class | Number of 2D classes: 10<br>Align filament classes vertically: true | 10 particle classes and 10 templates |
| 20 | Select 2D | Output #19 | - | TMV: 4 classes<br>- 4,972 segments |
| 21 | TMV *ab-initio* reconstruction | Output of #21 | - | Initial model |
| 22 | Helical refinement<br>TMV | Selected classes,<br>*ab-initio* model | Helical twist:<br>22.04<br>Helical rise: 1.408<br>High-resolution<br>noise substitution:<br>true | Final map |

## Results and Discussion

Four components were chosen for the EM ladder: ApoF, β-gal, PP7 and TMV. ApoF is an octahedral protein cage and a widely used specimen for characterization of the resolution limit of cryoEM microscopes. Mouse ApoF has yielded the highest available resolution from cryoEM single-particle analysis. Commercially available horse spleen ApoF can be used resulting in ∼2 Å reconstructions ^8,16^. However, ApoF is notoriously difficult to reconstruct from poor-quality data acquired from thick ice, charge-coupled detectors (CCDs), or low-voltage microscopes ^17–19^, so β-gal in this EM ladder mix serves as a low-symmetry (D2) standard. It is commercially available and has been used extensively as a high-resolution test specimen. Prior to vitrification, β-gal can be incubated with various ligands, such as PETG, for additional stability ^6^. β-gal can be replaced in this mix with aldolase, conalbumin or any other low-symmetry small protein, although particle concentration and ratio of the mix may differ. PP7 VLPs are ∼22 nm icosahedral particles that can be reconstructed in a broad resolution range - from 30 Å to 2.5 Å. PP7 particles are twice the diameter of ApoF and can serve as an alternative for ApoF as a pixel size calibration standard. PP7 VLPs can be replaced with other large particles such as AAV, Q-beta, proteasome or GroEL. Finally, TMV has been used extensively as a resolution test specimen. It can be effectively used as another alternative for pixel size calibration and for processing as it provides a good starting point in learning helical processing. TMV can be replaced with similar helical viruses, microtubules or other helically-symmetric elongated constructs.

3,996 micrographs of the EM ladder were acquired, and each micrograph was manually selected for containing at least 3 species in the field of view, good ice and little to no gold included in the image. All four species can be easily identified in a micrograph - ApoF “donuts”, small spindle-shaped β-gal, large hexagons of PP7 VLPs and “train tracks” of TMV (Figure 1A). Processing these data also provides clear and easily interpretable results from particle picking, through 2D classification and initial model generation, to 3D refinement (Figure 1B). Despite the different sizes and symmetry types present in this dataset, initial processing can be done in only 22 cryoSPARC jobs following a one-shot processing strategy, where a single box size is used for all particles (Table 1, Figure S1). This results in sub-3 Å reconstructions for two species - the most abundant ApoF and helical TMV. Particle picking, box size, and classification and refinement strategies can be optimized from this point (Figure S2-S5). Final reconstructions for this multi-species dataset resulted in a GSFSC resolution of 2.47 Å for ApoF, 2.74 Å for β-gal, 3.37 Å for PP7 and 2.46 Å for TMV (Figure 2).

**Figure 1.**
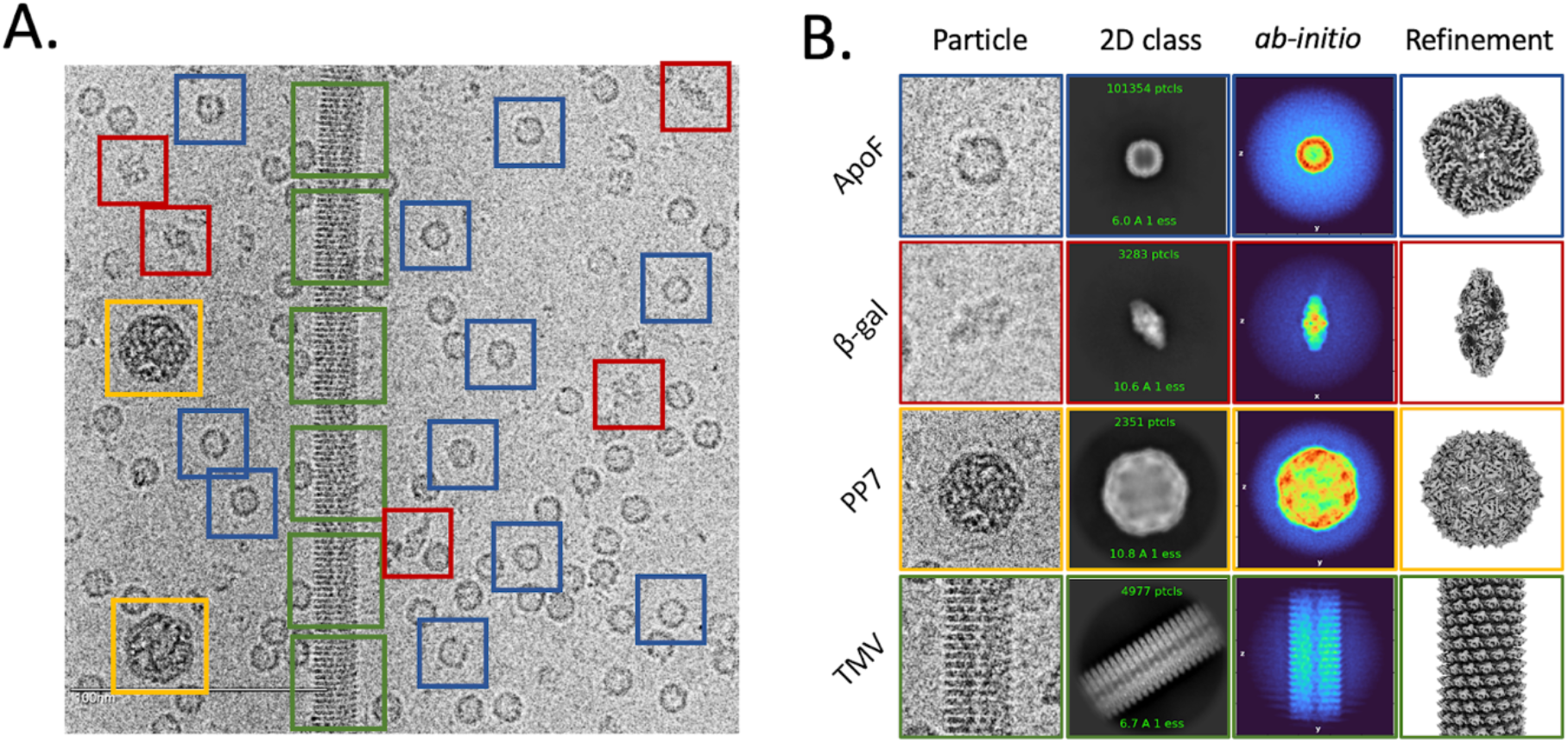
Multi-species dataset overview. A) An exemplar micrograph with four types of particles boxed out. B) Typical processing results: individual extracted particles, 2D class averages, x-y slice through the center of *ab-initio* reconstruction, 3D map visualized in ChimeraX.

**Figure 2.**
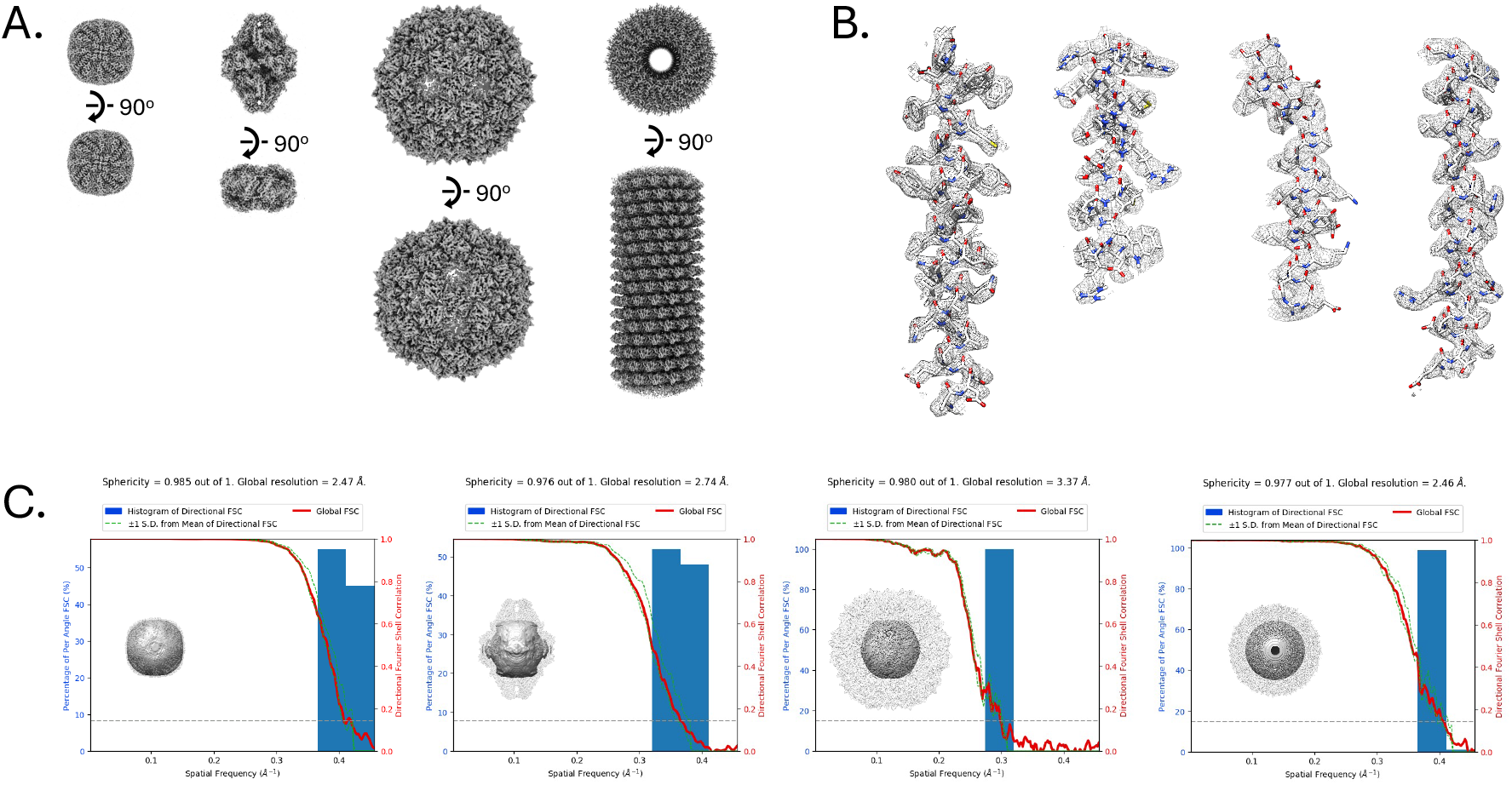
CryoEM reconstructions from multi-species dataset. A) Isosurface representations of *(from left to right)*: ApoF, β-gal, TMV, and PP7. B) PDB model fit to map in mesh of *(from left to right)*: ApoF 1FHA chain A: 14-42, β-gal 6X1Q chain A: 429-448, TMV 6R7M chain A: 107-136, and PP7 1DWN chain A: 96-121. C) Histogram and directional FSC plot with sphericity representation and transparent isosurface view *(from left to right)*: ApoF at a global resolution of 2.47 Å with 164,200 particles and sphericity of 0.985, β-gal at a global resolution of 2.74 Å with 37,617 particles and sphericity of 0.976, TMV at a global resolution of 2.46 Å with 12,038 particles and sphericity of 0.977, and PP7 at a global resolution of 3.37 Å with 1,803 particles and sphericity of 0.980.

After negative stain screening of the initial mix, the concentration of ApoF was halved due to it being the dominant protein in the micrographs (Figure 3). Each mix was subsequently changed based on the amount of each protein seen during screening. The decision to switch to cryo for screening was made after observation that a mix that had a good distribution of each protein in negative stain did not have the same distribution in cryo. During cryo screening, broken ends of thyroglobuin (ThG) were visible in the background of the micrographs. ThG requires detergent, such as CHAPS, to remain fully intact in cryo. For further mixes, ThG was switched to β-gal in order for all proteins in the mix to be fully reconstructed without adding detergent to the buffer.

**Figure 3.**
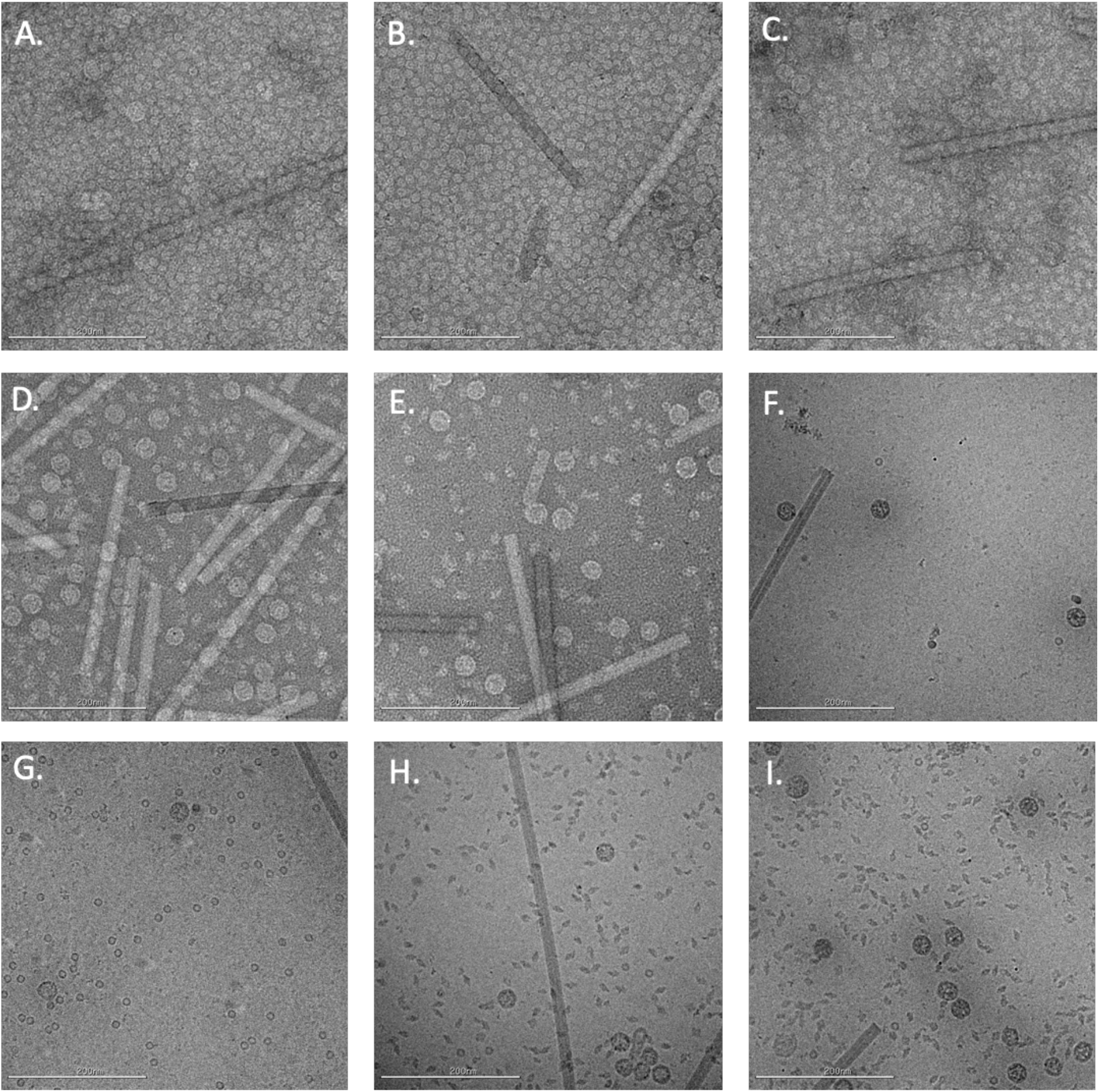
Representative images from trials of multi-protein mixtures. A through E: negative stain micrographs of mixes 1 through 5. F through I: cryo micrographs of mixes 6 through 9. A=mix 1, B=mix 2, C=mix 3, D=mix 4, E=mix 5, F=mix 6, G=mix 7, H=mix 8, I=mix 9. Exact mix compositions are shown in Table 2.

Although ApoF has become a widely used benchmarking standard for high resolution testing of all transmission electron microscopes ^20–22^, it still presents a challenge when processing. Due to the similar size of β-gal and ApoF, having both present within a micrograph is useful to test several particle pickers for their ability to differentiate between the two species, while also being able to benchmark the microscope. In contrast to ApoF, VLPs are larger, simple to pick, and can be used for low resolution testing as well as calibrating pixel sizes. In addition to the single particle processing practice, including TMV in the mix presents the opportunity to learn or improve the ability to do helical processing. TMV has been widely studied, so knowing the pitch, rise and twist is helpful whether you are processing from scratch and needing to confirm parameters, or if you are just trying to learn the different steps involved in helical processing versus single particle processing.

**Table 2.** Protein composition mix trials with representative images in Figure 3. Concentrations are in mg/ml.

| Mix (#) | TMV | PP7 | ApoF | ThG | β-gal |
| --- | --- | --- | --- | --- | --- |
| 1 | 0.17 (0.0425) | 0.05 (0.0125) | 0.08 (0.02) | 0.1 (0.025) |  |
| 2 | 0.17 (0.0425) | 0.05 (0.0125) | 0.004 (0.001) | 0.1 (0.025) |  |
| 3 | 0.17 (0.0425) | 0.05 (0.0125) | 0.00008 (0.00002) | 1 (0.25) |  |
| 4 | 0.17 (0.0425) | 0.05 (0.0125) | 0.00008 (0.00002) | 0.1 (0.025) |  |
| 5 | 0.17 (0.0425) | 0.05 (0.0125) | 0.0004 (0.0001) | 0.1 (0.025) |  |
| 6 | 0.17 (0.0425) | 0.05 (0.0125) | 0.04 (0.01) | 0.1 (0.025) |  |
| 7 | 0.17 (0.0425) | 0.05 (0.0125) | 0.08 (0.02) | 0.1 (0.025) |  |
| 8 | 0.17 (0.0425) | 0.05 (0.0125) | 0.08 (0.02) |  | 1 (0.25) |
| 9 | 0.17 (0.0425) | 0.1 (0.025) | 0.08 (0.02) |  | 1 (0.25) |
| 10 | 0.17 (0.0425) | 0.1 (0.025) | 0.16 (0.04) |  | 1 (0.25) |

As new practitioners adopt cryoEM workflows in their research programs, they need guidelines to not only ensure that their pipelines are compatible with their experimental requirements but also to be confident that their instrumentation can yield reliable and accurate results. Within a single experiment using a cryoEM calibration standard, researchers can validate a particular imaging condition or generate training datasets that can be used for benchmarking and cryoEM education.

## Conclusions

The introduction of dedicated cryoEM calibration standards extends far beyond benchmarking instrumentation by providing baseline control experiments for researchers to standardize their entire workflow for use in biophysical and structural biology research. The EM ladder can be used for high-resolution reconstructions but ensure the quality and reproducibility of data. Furthermore, this standardization endeavor extends to imaging settings, offering a reliable foundation for electron microscopes to consistently cater to a diverse array of biological samples. As cryoEM methodologies continue to evolve, control experiments form a necessary foundation of consistency and precision for integrative analysis. Taken together, the cryoEM calibration standard concept facilitates cross-disciplinary structural biology methodologies that rely on the reproducibility of cryoEM data.

## Acknowledgements

This work was performed at the Simons Electron Microscopy Center located at the New York Structural Biology Center, supported by grants from the Simons Foundation (SF349247), NYSTAR, NY State Assembly, and the NIH National Institute of General Medical Sciences (GM103310) with additional support from U24 GM129539. We thank Louise Fairall and Christos Savvy from the Protex facility at the University of Leicester for ferritin human H chain, and Ruben Diaz-Avalos from the La Jolla Institute for Immunology for the tobacco mosaic virus samples.

## Data availability

The cryoEM maps of all four components processed following the workflows described here were deposited to the Electron Microscopy Data Bank (EMDB) with accession codes: Human Apoferritin: EMD-41923, β-galactosidase: EMD-41919, PP7 virus-like particles: EMD-41917, and Tobacco mosaic virus: EMD-41924. Electron micrographs were made available at the Electron Microscopy Image Archive (EMPIAR) with accession code EMPIAR-11693. VLP aliquots may be provided upon request.

