## Supplemental information for "Multi-species cryoEM calibration and workflow verification standard"

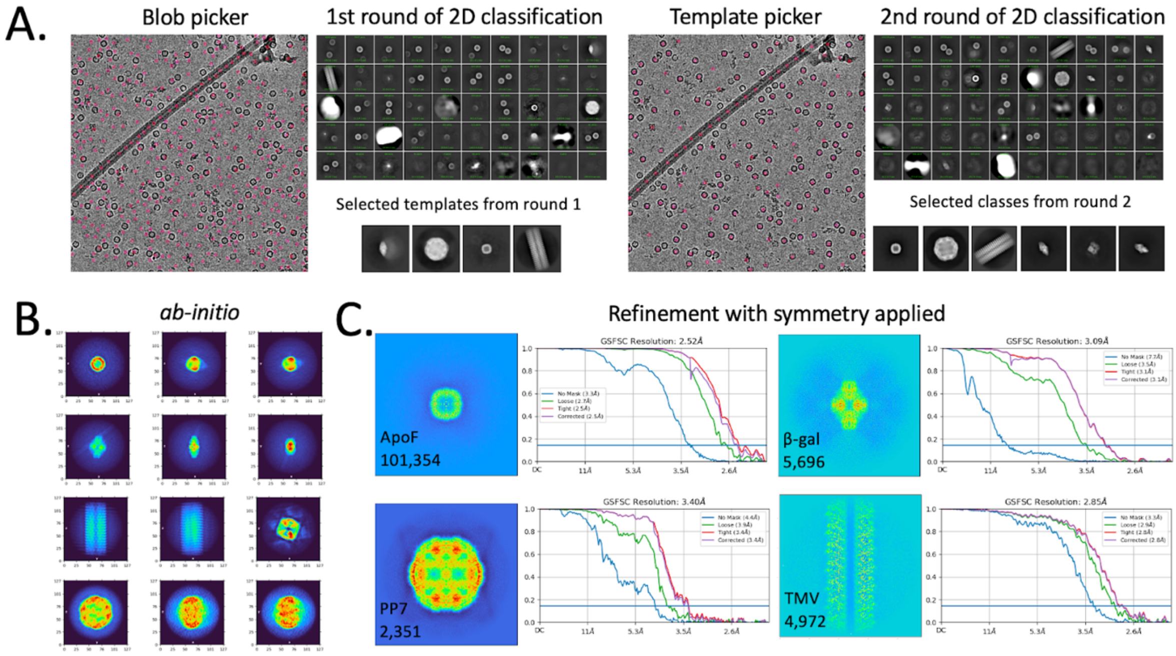


**Figure S1. One-shot processing strategy in cryoSPARC.** A) Diagnostic images from two rounds of particle picking and 2D classification. First blob picking was used, and templates were selected for the second round of picking using template picker. B) Diagnostic images from *ab-initio* reconstruction of particles from selected round two 2D classes. C) Output of 3D refinement jobs for all four species. Step-by-step information on jobs and settings for one-shot processing in cryoSPARC is described in Table 1.

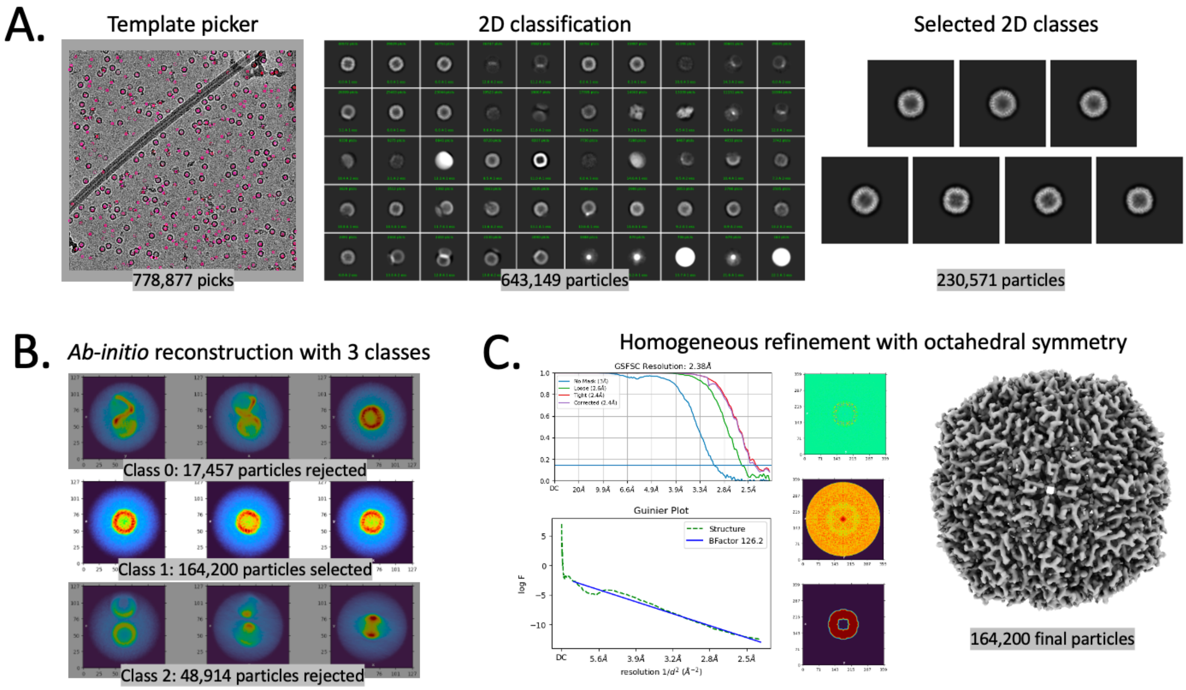


**Figure S2. ApoF processing.** ApoF template from one-shot processing was used for template picking

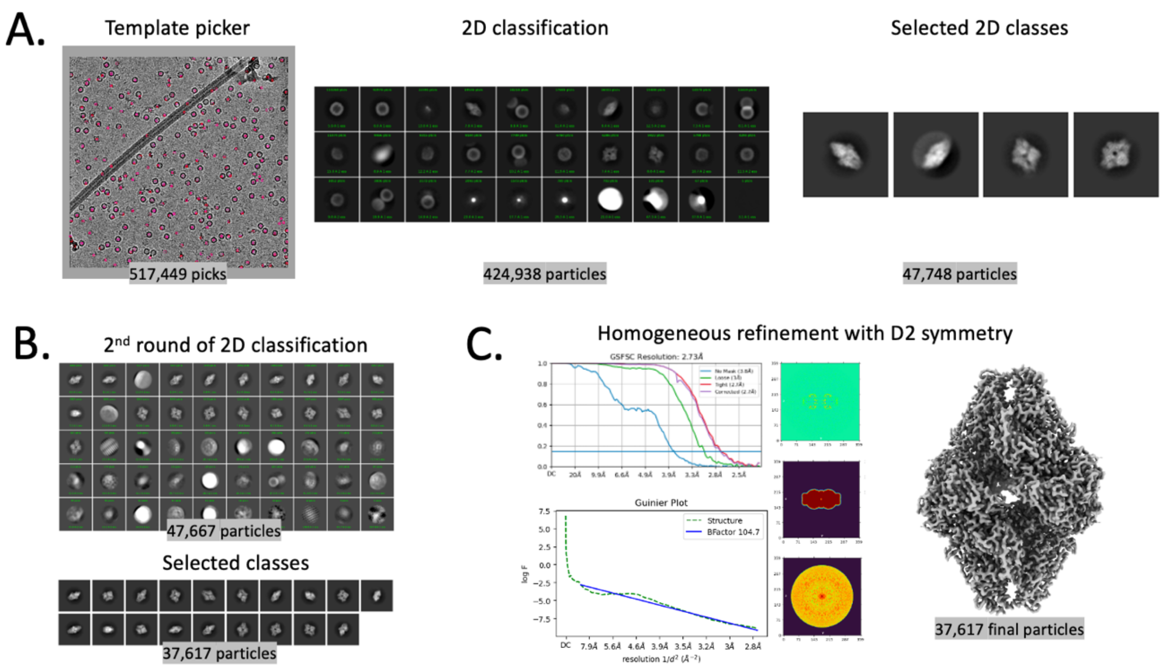


**Figure S3. β-gal processing.**

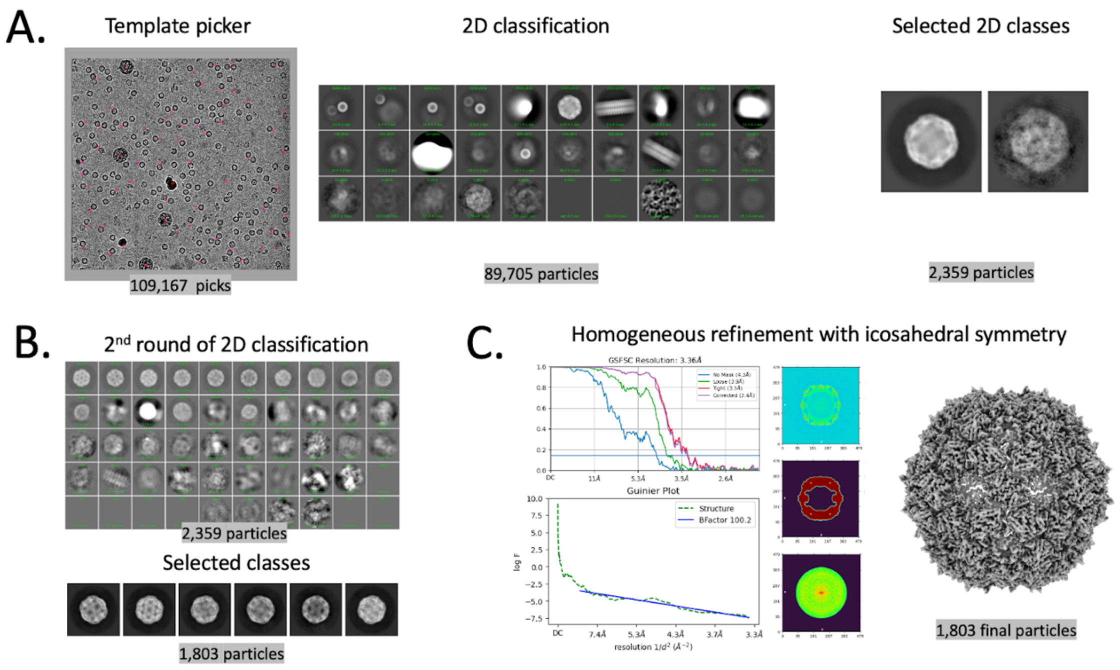


**Figure S4. VLP processing.**

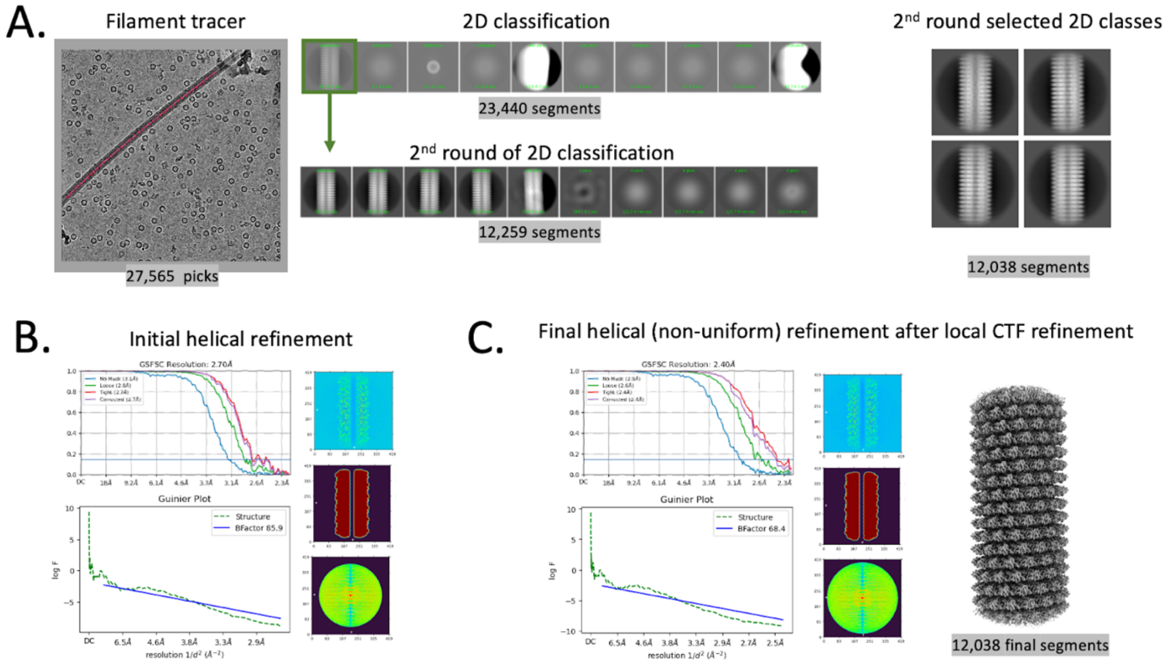


**Figure S5. TMV processing.**
